# Grape extract and resveratrol mitigate sleep fragmentation, Aβ accumulation, and abnormal neuronal excitability in a *Drosophila* model of Alzheimer’s disease

**DOI:** 10.1101/2024.07.14.603461

**Authors:** Ziqi Yu, Yangkun Xu, Yong Ping

**Affiliations:** Bio-X Institutes, Key Laboratory for the Genetics of Developmental and Neuropsychiatric Disorders (Ministry of Education), Shanghai Jiao Tong University, Shanghai 200240, China; Tandon School of Engineering, New York University, Brooklyn, NY, 11201, USA

## Abstract

Consumption of red wine and grape extracts may offer a range of health benefits, largely attributable to the grapes’ rich content of vitamins, fiber, and antioxidant compounds, such as polyphenols. To determine if resveratrol (RES) present in grape extracts is responsible for these benefits, we conducted a study on the effects of red grape skin extract (GSKE), seed extract (GSEE), and RES on sleep patterns, amyloid-beta (Aβ) deposition, neuronal excitability, and lifespan in a *Drosophila* model expressing Aβ42. Aβ42 flies experienced significant sleep fragmentation at night, yet their overall sleep duration was unaffected. Dietary GSKE significantly enhanced sleep duration and mitigated sleep fragmentation in these flies, whereas GSEE only increased the duration of sleep bouts during the day. RES demonstrated a similar effect, albeit to a lesser extent compared to GSKE. All three dietary interventions led to a reduction in Aβ42 levels and an extension of the lifespan in Aβ42 flies, with GSEE showing the least pronounced effects. Furthermore, GSEE and RES were able to reverse the hyperexcitability of mushroom body neurons (MBNs) caused by Aβ42 expression. These results suggest that GSKE and RES are potent promoters of sleep and have the potential to ameliorate sleep disturbances. Additionally, the study highlights that other bioactive component in GSKE, beyond RES, may contribute to its diverse pharmacological activities, which could differ from those of GSEE or RES alone. This underscores the multifaceted nature of grape extracts and their potential therapeutic applications in addressing sleep disorders and neurodegenerative conditions associated with Aβ deposition.

## Introduction

Grapes are commonly consumed all over the world. The health benefits of red wine consumption have been widely reported, from protecting against heart diseases, cancers to neurological diseases [1, 2]. Hundreds of compounds were found in red wine, including sugars, organic acids, alcohols, minerals and polyphenols, and most of health benefits were identified largely attributable to polyphenolic compounds [3]. Phenolic compounds were classified into flavonoids and non-flavonoids. Dietary foods and wines contain a diversity of flavonoids and non-flavonoids, so it is difficult to determine the function of specific compounds, and their specific contribution to the beneficial health effects *in vivo* [4]. Although bioactive compound contents could be dependent on the variety, geographical location and health of the grape, the compounds in grape skin extract (GSKE) and seed extract (GSEE) could also be quantitatively different [5]. Basically, non-flavonoids (such as stilbenes) were mainly found in GSKE, while flavonoids (such as proanthocyanins) were more abundant in GSEE [6, 7]. In particular, a group of polyphenolic compounds named stilbenes (belonging to non-flavonoids), especially resveratrol (RES), contribute to antioxidant and anti-inflammatory responses [8]. Since GSKE and GSEE contain different bioactive compounds as stated above, they may exert their protections against various diseases through different pathways [9].

RES has also been suggested as a putative anti-aging bioactive compound for prevention of Alzheimer’s disease (AD) [10-13]. Some studies also suggested that grape extract and RES can modulate or treat AD pathology [14]. However, the potential differences between health benefits mediated by grape extract and RES in treating AD is unclear. *Drosophila* has been established as a valuable model for pathological study of AD and for sleep/wake analyses [15-18]. Impaired sleep patterns, such as sleep fragmentation, and neuronal activities are common in AD patients and animal models [19-21]. We have previously showed that RES reverses sleep defects in a *Drosophila* AD model [22], but whether grape extract can restore sleep and aberrant neuronal activities is unclear. In this study we intend to compared the health benefits by dietary of RES, GSKE and GSEE in AD and control *Drosophila* models, and these data could help evaluate the potential contribution of RES among dozens of major bioactive compounds found in grape extract.

## Results

### GSKE and RES, but not GSEE, alleviate sleep fragmentation

*Drosophila* models expressing human Aβ42 provide great opportunities for investigating AD-related neuropathology [23]. We have previously showed that *Drosophila* AD models exhibited sleep disorders, including sleep fragmentation and impaired sleep homeostasis [21, 24]. In this work, we first tested possible effects of constitutively expressing Aβ42 in neurons (*Elav-Gal4>UAS-Aβ42*, Aβ42) on sleep in flies. We found that general sleep time was unaltered compared to control lines, whereas bout number was significantly increased accompanied with reduced bout duration during night-time, but not for day-time (Figure 1A-1D). These data indicate that Aβ42 flies exhibit sleep fragmentation during night-time. To tested potential effects of GSKE, GSEE and RES on sleep patterns in Aβ42 flies, we made GSKE- and GSEE-supplemental fly food by adding 8% w/v of GSKE and GSEE powder to the normal food, according to previous reports [25, 26]. Dietary GSKE significantly increased day- and night-time sleep in Aβ42 flies, but not for GSEE (Figure 1A and 1B). Day-time sleep was also increased by dietary RES (250 μM/L). More importantly, GSKE and RES significantly reduced night-time bout number in Aβ42 flies, accompanied with increased bout duration (Figure 1C and 1D). Surprisingly, GSEE showed no effects on both bout number and duration (Figure 1C and 1D). These data indicate that distinct regulation of sleep patterns by GSKE and GSEE may attribute to containing distinct polyphenol contents. Specifically, RES, relatively much higher in GSKE, is a potent sleep promoter and could be largely responsible for alleviating sleep fragmentation by GSKE in Aβ42 flies.

**Figure 1.**
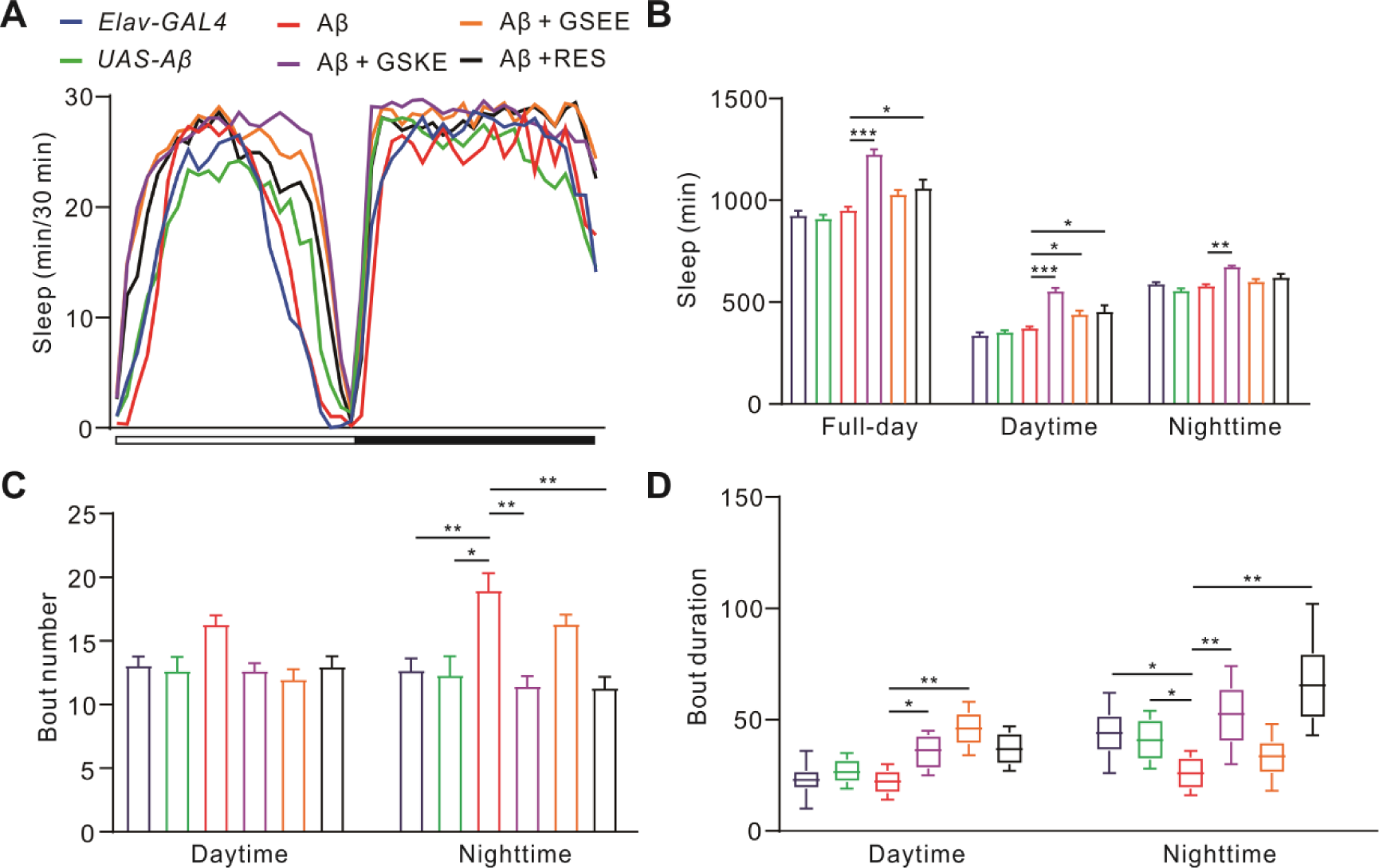
Grape extract reduces sleep fragmentation in Aβ42 flies. (A) Conventional sleep curves of controls and experimental lines, including *Elav-Gal4, UAS-Aβ42, Elav-Gal4>UAS-Aβ42* (Aβ42), Aβ42 + GSKE, Aβ42 + GSEE, Aβ42 + RES. All the flies were kept under 12h/12h LD cycles, aged 15-17 days. n = 30-42 for each group. (B) Total sleep time (full-day, 24 h) and sleep during daytime (12 h) and nighttime (12h) in control and experimental lines as shown in (A). (C) Quantification of mean sleep bout number during daytime and nighttime in the lines as shown in (A). (D) Box plots for daytime and nighttime sleep bout duration in the lines as shown in (A). The line inside the box indicates the median; the upper and lower box limits the 75% and 25% quantiles; vertical lines above and below the box represent the 95% and 5% quantiles. Data are presented as the mean ± SEM. \**P* <0.05, \*\**P* <0.01, \*\*\**P* <0.001. One-way ANOVA followed by post hoc Tukey was used to test hypotheses involving multiple groups. The Mann-Whitney U test was applied for the comparison of non-parametrically distributed data in (D).

### Grape extract reduces Aβ42 deposits

We have shown that GSKE alleviates sleep fragmentation in Aβ42 flies, which were also shown to exhibit Aβ accumulation in mushroom body and related pathology, including dramatic neuronal loss [17]. We asked whether grape extract could help Aβ clearance in the fly model. Thus, we quantified Aβ42 level by 6E10 antibody, and our data showed accumulating Aβ42 deposits in Aβ42 flies aged 15 days, and much more stronger in flies aged 25 days (Figure 2A and 2B). No Aβ42 signals were found in control lines. Also, we noticed obvious block holes (probably neuronal loss) in older flies aged 25 days caused by Aβ-related neuropathology (Figure 2A). Interestingly, either dietary GSKE, GSEE or RES significantly reduced Aβ42 levels at different ages (Figure 2B).

**Figure 2.**
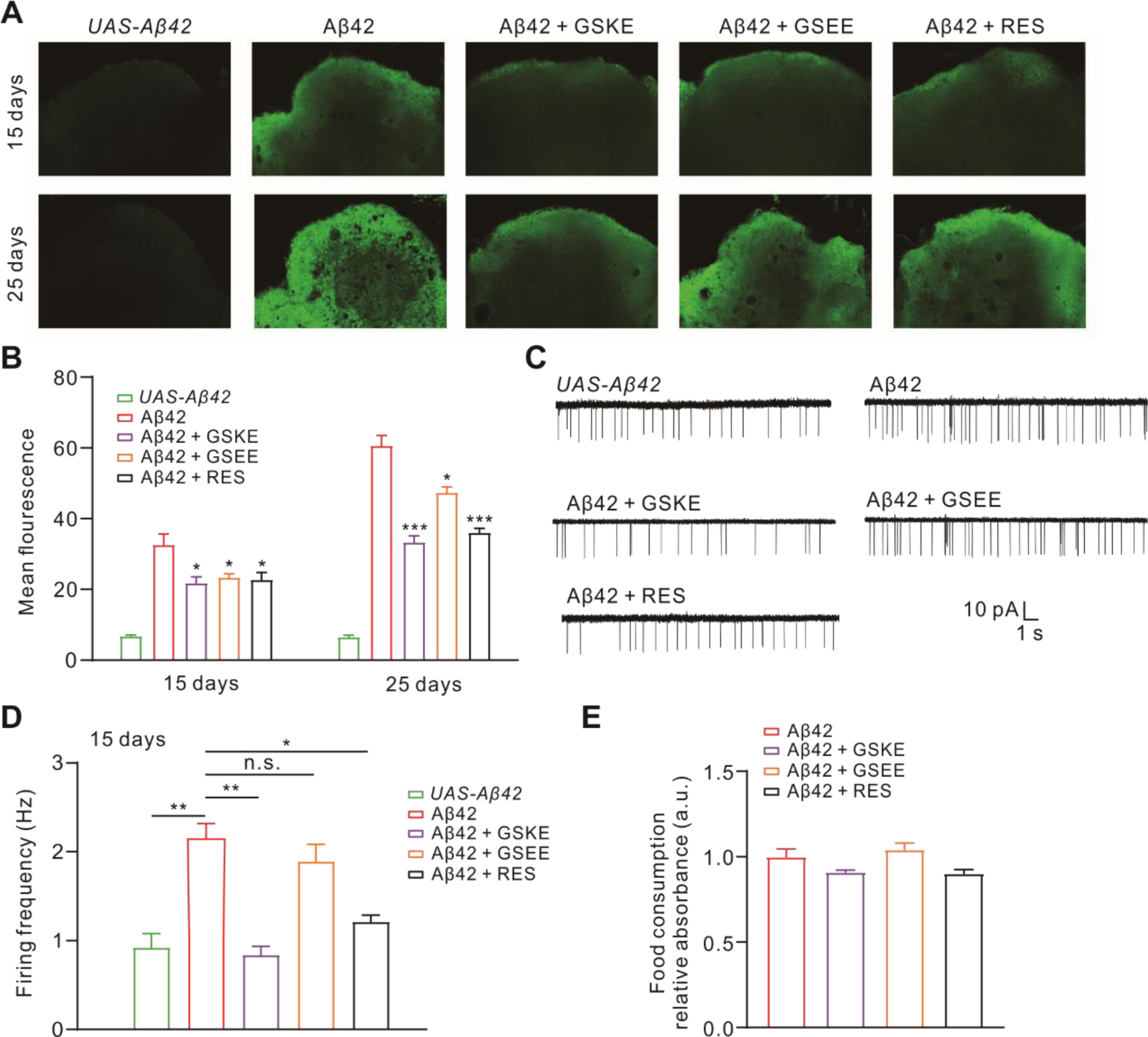
Grape extract reduces Aβ42 aggregates and neuronal excitability. (A) Representative confocal images of mushroom body area stained by 6E10 (Aβ42) antibody (red) in the indicated genotypes and treatments: *UAS-Aβ42, Elav-Gal4>UAS-Aβ42* (Aβ42), Aβ42 + GSKE, Aβ42 + GSEE, Aβ42 + RES. (B) Quantification of relative Aβ42 fluorescence (measured as grey values) in the indicated genotypes as described above. n = 10-12 brains for each group. (C) Typical firing patterns were recorded from MBNs of indicated genotypes and treatments, including *UAS-Aβ42, Elav-Gal4>UAS-Aβ42* (Aβ42), Aβ42 + GSKE, Aβ42 + GSEE, Aβ42 + RES. (D) Quantification of firing frequency in the cells as shown in (C). n = 12-15 cells for each group. (E) Quantification of food consumption of the Aβ42 flies under normal food, or additional with skin, seed or RES. We measured by using a colorimetric estimation after the treatment with non-absorbed blue dye. n = 5 times for each group Data are presented as the mean ± SEM. \**P* <0.05, \*\**P* <0.01, \*\*\**P* <0.001. One-way ANOVA followed by post hoc Tukey was used to test hypotheses involving multiple groups.

We have previously showed neuronal hyperexcitability in the *Drosophila* Aβ42 model [17, 27], and then we wondered if grape extract and RES attenuate hyperexcitability. we monitored spontaneous activity of mushroom body neurons (MBNs) labeled by a *GFP*.*S65T*.*T10* transgene, which promote GFP expression only in MB neurons, regardless of the GAL4 driver present. We applied cell-attached voltage-clamp of neuronal activity without changing the cytoplasmic milieu to record neuronal firings from the line *Elav-Gal4>UAS-Aβ42;UAS-GFP*.*S65T*.*T10*. Our data showed that GSKE and RES significantly reduced firing rate in MBNs of Aβ42 flies, while GSEE had no effects on firing rate (Figure 2C and 2D).

We also tested whether the three supplements affect food intake in Aβ42 flies. No effect was observed on food intake by GSKE, GSEE or RES (Figure 2E), measured by quantifying a food dye added to food as indicated, demonstrating that food intake was unaltered by dietary supplements.

### Grape extract extends lifespan in Aβ42 flies

Since a shortened lifespan has been observed in *Drosophila* Alzheimer’s models, we investigated whether constitutive expression Aβ42 can induce premature death. Our data showed that the survivorship of Aβ42 flies was greatly impaired. Aβ42 flies showed a median age of ∼34 days, significantly shorter than then control lines (∼50 and ∼53 days for *Elav-Gal4* and *UAS-Aβ42* lines, respectively) (Figure 3). Interestingly, either GSKE, GSEE or RES supplement resulted in a significantly increase in lifespan of Aβ42 flies (median age at ∼51, ∼39 and ∼47 days for GSKE, GSEE and RES supplements, receptively) (Figure 3). These data support the notion that grape extract and RES reduce premature death in Aβ42 flies.

**Figure 3.**
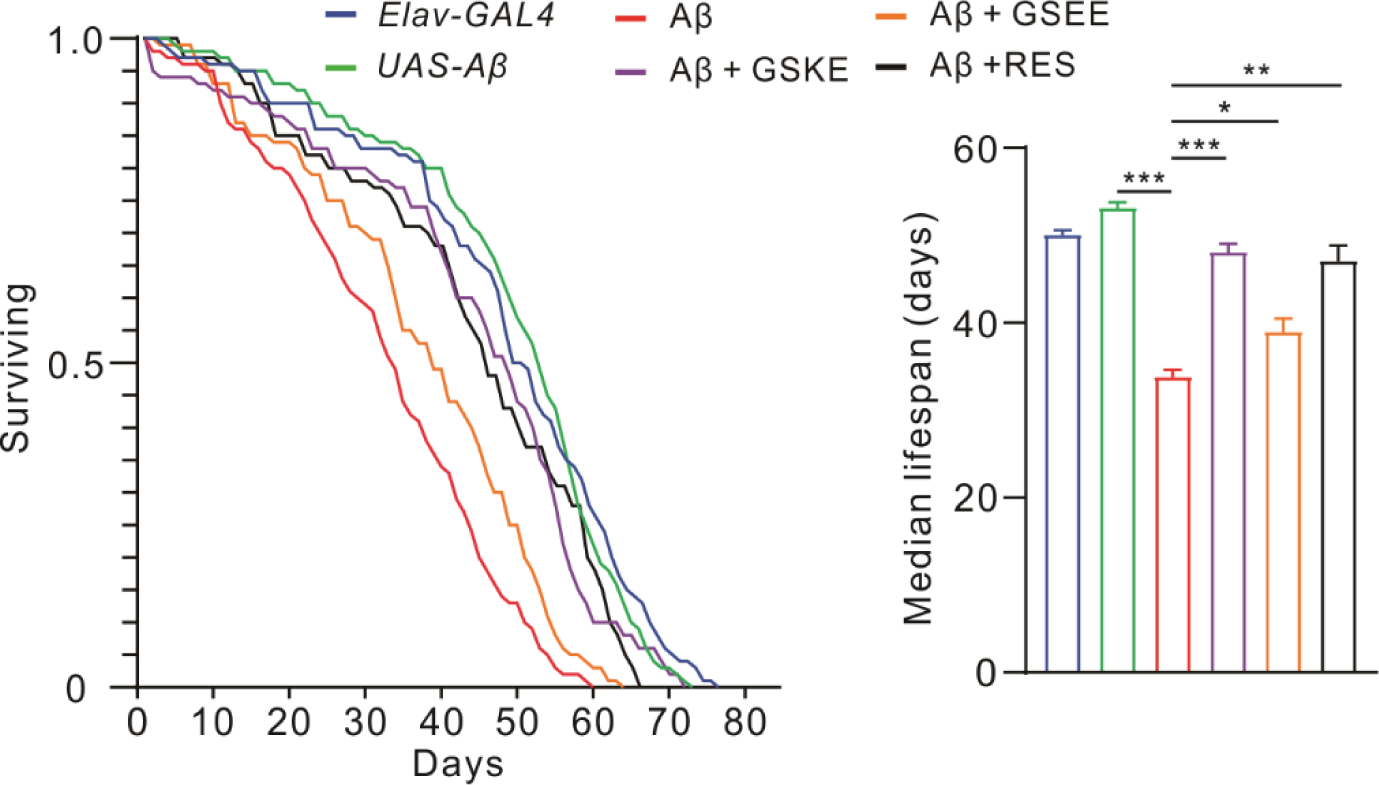
Effects of GSKE, GSEE and RES on the lifespan of Aβ42 flies. Lifespan of control and Aβ42 flies fed with normal food, and Aβ42 flies raised on normal food supplemented with GSKE, GSEE or RES (left panel). Quantification of the median lifespan of control and Aβ42 flies subjected to the various treatment conditions as shown in the left panel (right panel). Data are presented as the mean ± SEM. \**P* <0.05, \*\**P* <0.01, \*\*\**P* <0.001. One-way ANOVA followed by post hoc Tukey was used to test hypotheses involving multiple groups.

## Discussion

AD is a common neurological disease with complex neuropathology. Attempts to treat the disease has largely failed and there are only a few medications available. At this stage, perhaps we may need to consider other alternative strategies, including bioactive compounds from plants. For example, various polyphenols, secondary metabolites of plants, has been shown to suppress Aβ aggregation and modulate AD and other neurodegenerative pathology [28-31]. Although the effects of grape extract on Aβ deposition, antioxidant and anti-inflammatory responses have been extensively investigated, their potential effects on sleep patterns and neuronal excitability are largely unclear. We have previously showed that RES promote sleep in a fly AD model, but whether RES containing grape extract can alleviate sleep disorders is unknown. So, in this study we investigated neuroprotective effects of GSKE, GSEE and RES in Aβ42 flies, with a focus on sleep regulation. We found that unlike GSKE and RES, GSEE exhibited little effects on regulating sleep patterns in Aβ42 flies.

Emerging data have shown a strong link between sleep/circadian disorders and Aβ42 accumulation/AD [32, 33]. Increased nighttime activity and fragmented sleep are hallmarks of breakdown sleep/wake cycle in AD [34]. Sleep disorders in AD animal models may vary from one to another, but generally they exhibited increased sleep fragmentation and reduced sleep time [19]. We have previously showed increased sleep fragmentation with shortened sleep time in temperature controlled Aβ42 expression model [21]. Consistent with these findings, we found increased sleep fragmentation in constitutive Aβ42 expression model. GSKE and RES supplements reversed sleep fragmentation, while surprisingly GSEE showed little effects on sleep in Aβ42 flies. We propose that it’s probably due to relatively higher non-flavonoids, especially RES in GSKE. But we cannot exclude other bioactive compounds in GSKE contributing to sleep regulation.

Improved sleep has been suggested to enhance Aβ clearance and reduce Aβ deposition [19, 21]. Indeed, we observed reduced Aβ levels by dietary GSKE and RES. Interestingly, we found that dietary GSEE also reduced Aβ levels, although the effect was much weaker then GSKE and RES. Similar findings were also observed in our lifespan assay, showing that in addition to GSKE and RES supplements, GSEE also increased lifespan of Aβ42 flies. These data suggested that sleep patterns, Aβ deposition and lifespan can be regulated by distinct bioactive compounds and GSEE could enhance Aβ42 clearance through anti-oxidation and/or anti-inflammation, etc. Further work is needed to demonstrate whether reversing sleep fragmentation could further enhance Aβ42 clearance in the model as we expected.

Continuous accumulation of Aβ and neuronal hyperexcitability are common at early stage of AD [35, 36]. Interestingly, RES was found to control neuronal excitability under pathological conditions, including Aβ treatment [37], seizures [38], hyperalgesia [39]. Consistent with our data, we found neuronal hyperexcitability in our AD model, and we further showed that RES and GSKE reverse neuronal hyperexcitability, but not for GSEE. It would be interesting to know the potential link between neuronal hyperexcitability and sleep fragmentation observed in AD models. A recent report showed aberrant neuronal excitability in hypothalamic neurons contributed to sleep disorders in a mice AD model [40], indicating the potential bidirectional regulations between neuronal activity and sleep in AD pathology.

Finally, several studies have suggested that bright light therapy and physical exercise training could help increase sleep quality in old adults or AD patients [41, 42]. Since discovering new treatment strategies that could improve sleep quality in AD is important, our study provides a new way, whereby using GSKE and RES supplements, that could possibly enhance sleep quality in sleep disordered animals. Further studies are needed for determining their effectiveness and underlying mechanisms in managing AD-related symptoms.

## Methods

### Fly stock

Flies were maintained at 25 °C with a 12-h light/dark (LD) cycle on standard fly food and male flies were used in the study. *Elav-Gal4* (458, Chr X) was obtained from the Bloomington *Drosophila* Stock Center. *UAS-Aβ42* was provided by Dr. Yi Zhong [23]. *UAS-GFP*.*S65T*.*T10* was generated by Kei Ito group [43]. Flies were outcrossed for more than five generations with the *w*^*1118*^ (5905) strain to standardize the background before behavioral and lifespan experiments.

### Grape extract and RES

Red grape skin extract (GSKE) and seed extract (GSEE) under study were obtained from Sanxin Biotech (Hubei, China). The powders were stored at room temperature in a dry place. We added GSKE or GSEE at 8% (v/w) to the fresh fly food after curing, just prior to cooling. RES was firstly dissolved in ethanol and then added to fresh food before cooling at final concentration of 200 μM.

### Sleep assay

Male flies aged 12-14 days were collected and loaded into 5 mm (diameter) x 65 mm (length) glass tubes containing standard food or with GSKE/GSEE/RES supplements at indicated concentrations. The locomotor activities of flies were measured by the *Drosophila* Activity Monitoring System (DAMS, Trikinetics, MA) in incubators at 25°C with a standard 12h/12h LD cycle. All the flies were allowed to acclimate for 3 days before data collection. Activity counts were recorded at 1 min intervals, and the data were analyzed via custom a newly generated R package, named SleepyFlyR (GitHub-SoapMou/SleepyFlyR: SleepyFlyR: an R package for sleep and activity analysis in *Drosophila*). Sleep was identified based on the previously-established criterion that a sleep bout was defined as no movement within at least 5 min.

### Immunostaining analysis

Brains of flies at indicated ages were dissected and fixed in 4 % paraformaldehyde for 15 min and then were washed for 10 min, three times. We then blocked the brains in 5 % goat serum for 2 h and stained with mouse anti-Aβ42 (6E10) (1:500, BioLegend, CA, USA) in PBST on a shaker at 4 °C overnight. Brains were then washed for 10 min, three times, and secondary antibody was goat anti-mouse (1:500) was added for 2 h at room temperature. We obtained the images on a Leica SP5 confocal microscope.

For Aβ42 fluorescence quantification, regions of interest (ROIs) were chosen in the mushroom body area, which was essential for learning and memory, as previously reported [17]. Grey values of fluorescence labeled by 6E10 were further determined by subtracting the background, which was chosen from non-immunostained areas.

### Lifespan assay

Newly eclosed males of indicated genotypes or supplements in food were collected and separated into different vials (12 flies per group, 10 groups per experiments, with total 120 males per experiments), containing normal food or indicated supplements. Food vials were changed every 3 days and the number of dead/surviving flies was counted every day.

### Food intake measurement

To measure potential effects of RES, skin and seed extracts on food intake in *Drosophila*, we starved the male flies, aged 7 days, for 10 hours with water only. Then the animals were transferred to food containing 2.5% (w/v) food dye (Erioglaucine blue disodium salt, Sigma-Aldrich) for 12 hours at 25 °C. Flies fed on normal food were used as controls. After feeding, fly bodies were then collected with heads removed in PBS, and washed for 3 times. 10 bodies were homogenized in 300 mL of cold PBS, and centrifuged at 10,000 rpm for 10 min. Supernatant was used for absorbance recording at 625 nm using spectrophotometer.

### Electrophysiology

We dissected brains of transgenic flies (*Elav-Gal4>UAS-Aβ42;UAS-GFP*.*S65T*.*T10*) aged 15 days in culture medium (Invitrogen) and then we exchanged culture medium to the recording external solution, containing 110 NaCl, 2 KCl, 6 MgCl_2_, 1 CaCl_2_,5 glucose, 20 NaHCO_3_, 2 NaH_2_PO_4_ (in mM). Solution was bubbled with carbogen (5% O_2_ and 95% CO_2_) throughout recordings, with pH 7.2. All recordings were performed at room temperature. For cell-attached recordings, recording pipettes were filled with internal solution (K-gluconate, 120; KCl, 20; Mg-ATP, 4; Na-GTP, 0.5; HEPEs, 10; EGTA, 1.1; MgCl_2_, 2; CaCl_2_, 0.1, in mM, with pH 7.2), with additional 200 mM BaCl_2_ to prevent perforated patch effect mediated by Kir channels in the attached membrane. Cell attached configuration was done by gentle suction via a syringe. The recordings were performed in voltage-clamp mode with holding at 0 mV. Firing number was quantified using MiniAnalysis software for at least 1 min.

## Acknowledgements

We thank the Bloomington Stock Center for fly stocks used in this study. We thank members of the laboratory for discussion and assistance. This work was funded by the grants from the Shanghai Rising-Star Program (19QA1404900).

## Author contributions

YP conceived the study; ZY, YX and YP designed the experiments; ZY, YX, and YP performed experiments and analysed data; YP and ZY wrote the paper.

## Data availability statement

All the data that support the findings have been provided with the manuscript, or deposited in GitHub as noted in the text. Additional information, images, analysis data are available from the corresponding author, upon reasonable request.

